# RFQAmodel: Random Forest Quality Assessment to identify a predicted protein structure in the correct fold

**DOI:** 10.1101/654293

**Authors:** Clare E. West, Saulo H. P. de Oliveira, Charlotte M. Deane

## Abstract

While template-free protein structure prediction protocols now produce good quality models for many targets, modelling failure remains common. For these methods to be useful it is important that users can both choose the best model from the hundreds to thousands of models that are commonly generated for a target, and determine whether this model is likely to be correct. We have developed Random Forest Quality Assessment (RFQAmodel), which assesses whether models produced by a protein structure prediction pipeline have the correct fold. RFQAmodel uses a combination of existing quality assessment scores with two predicted contact map alignment scores. These alignment scores are able to identify correct models for targets that are not otherwise captured. Our classifier was trained on a large set of protein domains that are structurally diverse and evenly balanced in terms of protein features known to have an effect on modelling success, and then tested on a second set of 244 protein domains with a similar spread of properties. When models for each target in this second set were ranked according to the RFQAmodel score, the highest-ranking model had a high-confidence RFQAmodel score for 67 modelling targets, of which 52 had the correct fold. At the other end of the scale RFQAmodel correctly predicted that for 59 targets the highest-ranked model was incorrect. In comparisons to other methods we found that RFQAmodel is better able to identify correct models for targets where only a few of the models are correct. We found that RFQAmodel achieved a similar performance on the model sets for CASP12 and CASP13 free-modelling targets. Finally, by iteratively generating models and running RFQAmodel until a model is produced that is predicted to be correct with high confidence, we demonstrate how such a protocol can be used to focus computational efforts on difficult modelling targets.

## Introduction

Template-free protein structure prediction protocols routinely produce hundreds to thousands of models for a given target [1]. Users need to be able to identify if a good model exists in this ensemble. The final step in a typical structure prediction pipeline is therefore to select a representative subset of five or fewer models as output [2]. This model selection step is critical, and the community’s ability to select good models is assessed as part of the Critical Assessment of protein Structure Prediction (CASP) experiments [3].

Protocols for model quality assessment can be divided into three classes: single-model methods, quasi-single model methods, and consensus methods [2]. Single-model methods calculate a score for each model independently, and this score does not take into account any of the other models generated for a particular target. The objective function optimised during protein structure prediction can usually be used as a single-model quality estimator, but better results have been reported if different scores are used for modelling and ranking [2]. Examples of single-model scores include ProQ3D [4] and the ROSETTA energy terms [5]. For quasi-single model methods, the score of a given model is calculated based on its relative score compared to a subset of all models (reference set) produced for the target, for example MQAPsingle [6]. Consensus methods, such as Pcons [7], perform pairwise comparison of the predicted structures to identify clusters of similar models or regions, and assume that structures with high consensus are more likely to be correct.

Predicted contacts derived from co-evolution analysis of multiple sequence alignments have been used as single or quasi-single model methods to improve model quality assessment (e.g. [7, 8]). Existing contact-based methods for quality assessment often consider the proportion of predicted contacts that are satisfied in each model (i.e. how many of the pairs of residues predicted to be in contact are within a certain threshold distance) [7]. ModFOLD6, a quasi-single model quality assessment method, includes a term describing the local agreement with predicted contacts for each residue in the model [9]. An alternative way to use predicted contact information is to align predicted contact maps for a particular target to the observed contacts maps of models. Contact map alignment has been used to select regions of models to be hybridised [10] or to perform protein threading [11]. Until now, contact map alignment has not been used for model quality assessment, but the principles that govern these techniques should also be applicable for quality assessment tasks.

In combination with recent advances in model quality due to better contact prediction techniques, improvements in model quality assessment have made template-free protein structure prediction more reliable (e.g. [7, 8]). The most recent CASP competition demonstrated remarkable progress in the field: the highest-performing method produced a model in the correct fold (TM-score ≥0.5) in the top five models for 23 of 32 free-modelling target domains, although performance decreases when considering only the top model. This level of predictive ability has driven efforts to perform large-scale modelling of significant numbers of protein families without a member of known structure [10, 12]. While these studies offer reliable topologies for many protein families, the recall of their quality assessment protocol remains low enough that some predictions with the correct topology may not be identified. Furthermore, such studies were limited by the computational expense of model generation, opting either to produce models for a subset of these families of unknown structure [10] or to produce a reduced number of models per target [12].

In this paper, we introduce RFQAmodel, a random forest quality assessment classifier developed to evaluate models produced by template-free protein structure prediction pipelines. The classifier combines existing quality assessment scores with predicted contact map alignment scores. Unlike most established quality assessment methods, RFQAmodel is trained to evaluate whether models are in the correct fold (TM-score ≥ 0.5) rather than estimating the absolute model quality. For each model, RFQAmodel outputs an estimated probability that the model is correct. This probability can be used to estimate whether the model is correct with high, medium, or low confidence, or if modelling is predicted to have failed.

We compiled Training and Validation sets each comprising 244 structurally diverse protein domains. We ensured that these sets were well-balanced in terms of protein length, number of effective sequences [7], SCOP class [13], and other properties that are known to have an effect on modelling success. We used our sequential protein structure prediction protocol SAINT2 [1] to generate 500 models for each of the 488 protein domains. Using the Training set, we show that predicted contact map alignment scores are as effective for ranking models as existing state-of-the-art quality assessment scores. Furthermore, the models ranked highly by these contact map alignment scores are different from those ranked highly by conventional scores. We incorporate several state-of-the-art quality assessment scores alongside contact map alignment scores into a random forest classifier, RFQAmodel, which classifies models as correct (i.e. in the correct topology) or incorrect, and outperforms the component quality assessment scores. Of the 244 targets in the Validation set, RFQAmodel predicts that the highest-ranking model may be correct for 185 targets, of which 86 are correct (out of a possible 142 for which at least one correct model was generated by SAINT2). The 185 are further split by RFQAmodel into those where the highest-ranking model is predicted to be correct with high confidence, 67 targets, of which 52 are correct. Of the 59 targets predicted to be modelling failures, 5 had at least one correct model, and none had a correct highest-ranking model. We demonstrate that similar results are achieved when applied to the server models submitted to CASP12 and CASP13. Finally, we demonstrate how RFQAmodel can be used to estimate when sufficient models have been generated for a particular target, enabling more efficient use of computational power.

## Materials and methods

### Training and Validation Sets

To construct our Training and Validation data sets, we used the mapping between Pfam [14] domains and PDB [15] structures as available on the EBI repository in February 2017. To represent each of these families, we selected the first protein chain listed for that family (SI Table 1).

We annotated each of the protein chains according to the 2.06 stable build of SCOPe [13]. If the protein chain selected to represent a Pfam family was not annotated in SCOPe, we tested all the remaining members of the family sequentially (as ordered on the mapping) to maximise the number of Pfam families with SCOPe annotations (SI Table 2 and SI Fig 1).

We excluded all families longer than 250 residues, and performed a culling and cleaning process (SI Section 2) that resulted in a data set of 488 structurally diverse protein domains (SI Table 3). The average length and number of effective sequences, *B*_eff_, as defined in [7] (see SI Section 3), of these domains were similar to those of the original PDB-mapped and SCOPe-annotated Pfam domain sets.

The 488 protein domains were divided into Training and Validation sets of equal size. For each SCOP class, we selected two domains at a time in order of increasing *B*_eff_ and randomly assigned one to the Training and the other to the Validation set. We used the *B*_eff_ of the multiple sequence alignments used for contact prediction. While this ensured that the sets have similar *B*_eff_ medians and have roughly the same number of protein domains for each SCOP class, the overall length and resolution distributions differed between sets (SI Fig 2). In particular, proteins in the Validation set with *B*_eff_ <100 tended to be longer than proteins on the Training set with *B*_eff_ <100, which suggests that the Validation set may be more challenging for protein structure prediction.

### Protein Structure Prediction

To produce models for all targets in our Training and Validation sets, we used our fragment-assembly protocol SAINT2 [1] (for details, see SI Section 4 and [1]) with the parameters given in the original publication, with the exception of secondary structure prediction. We used DeepCNF Q8 to predict secondary structure, as DeepCNF Q8 had a slightly higher precision for targets with large *B*_eff_ values, and results in marginal improvements in fragments with predominantly loop secondary structure (see SI Section 4.1).

In order for SAINT2 to produce the best possible model, the optimal number of models to generate is 10,000 [1]. However, for the purpose of developing a quality assessment protocol, we estimated that only 500 models were required to produce correct models for a sizeable number of targets (see SI Section 5).

We used SAINT2 to produce 500 models for each target in our Training and Validation sets. We assessed the number of modelling successes - targets for which at least one correct model (TM-score ≥ 0.5 [16]). was produced - as well as the TM-score of the best model produced for each target.

### CASP12 and CASP13 Test Sets

To test our classifier on models produced by methods other than SAINT2, and to compare its performance to other quality assessment methods, we used the stage2 server models used in the blind test of model quality assessment methods at CASP12 and CASP13. These consist of the 150 top-ranking server models submitted for 60 targets each for CASP12 and CASP13 targets. The models, model quality predictions, and model quality evaluations were accessed from the CASP website (http://www.predictioncenter.org/downlo_adarea/). This resulted in a total of 17,976 models for 120 targets. The lengths of the target structures range from 41 to 863 residues, with an average length of 289 residues.

### Model Validation

To assess the quality of the models produced by SAINT2, we used TM-align to calculate TM-score [16]. We consider all models with a TM-score ≥ 0.5 to be in the correct topology [17].

### Classification Features

For model classification, we used a set of 58 features, which can be divided into three groups: target-specific (3), model-specific (12), and ensemble-specific (43). The target-specific features are calculated from the target’s sequence, and are common to all models produced for that target. The model-specific features are calculated for each model, and include five existing single-model quality assessment scores, a consensus method quality assessment score, two scores based on the predicted contacts, and three predicted contact map alignment scores. The ensemble-specific features are summary statistics (maximum, median, minimum, and spread) of our model-specific features calculated across all models produced for each target. For all methods we used SAINT2 models and the predicted contacts generated by metaPSICOV. We note that many of the assessment scores used were not originally trained using these inputs, so their performance may be worse than expected.

#### Target-specific features (3)

The domain length, the *B*_eff_, and the total number of predicted contacts output by metaPSICOV with a score greater than 0.5.

#### Single-model quality assessment scores (5)

The final modelling score output by SAINT2, and the global score output by ProQ3D and component scores ProQ2D, ProQRosFAD and ProQRosCenD [4]. ProQRosCenD and ProQRosFAD are based on the Rosetta centroid and full atom [5] energy functions, respectively, which were calculated on relaxed models with repacked side chains. Relaxation was carried out using the *ab initio* relax protocol of Rosetta 3.7 as described in [4]. For ranking models, we have additionally considered the SAINT2 score without its contact component (SAINT2 Raw); this was not included as a feature in the random forest classifier.

#### Consensus quality assessment score (2)

We used the global score output by Pcons [18] with standard parameters. We also include PcombC [12], a weighted sum of three features: the ProQ3D global score, the Pcons consensus score, and the proportion of predicted contacts present in the model (positive predictive value, PPV).

#### Contact-based features (2)

The contact component of the SAINT2 score (see [1] for more details) and the proportion of satisfied predicted contacts (positive predictive value, PPV). Here, we considered a predicted contact to be a satisfied if the C-*β* atoms (C-*α* in the case of glycine) of the two residues predicted to be in contact were less than 8Å apart in the model output by SAINT2.

#### Predicted contact map alignment scores (3)

We used BioPython [19] to calculate an observed contact map for each model, with an 8Å distance cut-off between residue C-*β* atoms (C-*α* in case of glycine). We aligned the observed contact maps to the predicted contact maps produced from the output of metaPSICOV stage1. Two methods of contact map alignment were tested: map align [10], and EigenTHREADER [11]. Map align uses a dynamic programming algorithm to perform local contact map alignment and identify consensus regions. We used as features the best hit score and the best hit length produced by map align. EigenTHREADER uses eigenvector decomposition and dynamic programming to align the principal eigenvectors of the two maps. For an ensemble of structures, EigenTHREADER assesses which of the models is most likely to be in the same fold as the one described by the reference predicted contact map, assigning a relative score per model. We used the score output by EigenTHREADER for each model as a feature.

#### Ensemble-specific features (43)

The maximum, minimum, median, and spread (the difference between the maximum and the median) of 10 of our 12 model-specific features, excluding map align’s hit length and the proportion and absolute number of satisfied predicted contacts, for which only the maximum value for each target is included. These features were calculated per target across all models.

## Results

### Modelling Results

Correct models were produced for 151 out of 244 protein domains in our Training set, and 145 out of 244 protein domains in our Validation set. This corresponds to around 60% of the targets in each set, in line with numbers reported previously [1].

When considering the modelling results according to three *B*_eff_ bins (SI Fig 7A), our results corroborate previous findings that modelling is more likely to succeed when more effective sequences are available [8]. We observe a modelling success rate of 46% for our Training set at *B*_eff_ values below 100, and a success rate of 69% for *B*_eff_ ≥ 1000. Across our three *B*_eff_ bins (SI Fig 7A), we observe comparable modelling results for the Training and Validation sets, both in terms of the success rate and the distribution of the TM-scores of the best model for each target, with marginally worse performance for Validation set targets with *B*_eff_ values below 100.

We also find that modelling success rates vary by SCOP class (SI Fig 7B). For our Training set, SAINT2 produced a correct model for 85% of all-*α* targets, 65% of *α*/*β* targets, 61% of *α*+*β* targets, and 30% of all-*β* targets. Comparable modelling success rates and distributions of TM-score of the best models were obtained for Training and Validation sets across all four SCOP classes.

Modelling success rates also depend on domain length (SI Fig 7C). We separated the targets in our Training and Validation sets into four domain length bins (50 to 99, 100 to 149, 150 to 199, 200 or more residues). As expected, modelling success rate decreases as targets increase in length. For our Training set, SAINT2 produced a correct model for 83% of the targets that were 50 to 99 residues-long, for 65% of targets that were 100 to 149 residues-long, for 41% of targets that were 150 to 199 residues-long, and 39% of targets longer than 200 residues. When considering the combined effect of *B*_eff_ and domain length, SAINT2 failed to produce a correct model for all targets longer than 200 residues with a *B*_eff_ < 100 (see SI Fig 8).

Given the effect of these three features on modelling success, it is important to ensure that Training and Validation sets have similar distributions of domain length, effective sequences, and SCOP classes. A validation set that is comprised of shorter targets, or that contains more targets with a high *B*_eff_, or a disproportionate number of *α*-helical targets may lead to overestimation of classification performance.

### Comparing Quality Assessment methods

To assess the usefulness of including predicted contact map alignment scores as features for model quality assessment, we compared these scores with ten other model quality estimators: three SAINT2 scores and seven existing quality assessment scores. We ranked the 500 models produced by SAINT2 for each of the 244 targets in our Training set according to each of these model quality scores. For each score, we assessed the number of targets for which the highest-ranking model was correct (TM-score ≥ 0.5). Given that the quality of models is dependent on the availability of a sufficient number of effective sequences (*B*_eff_), we stratified this comparison across three *B*_eff_ bins (Fig 1).

**Fig 1.**
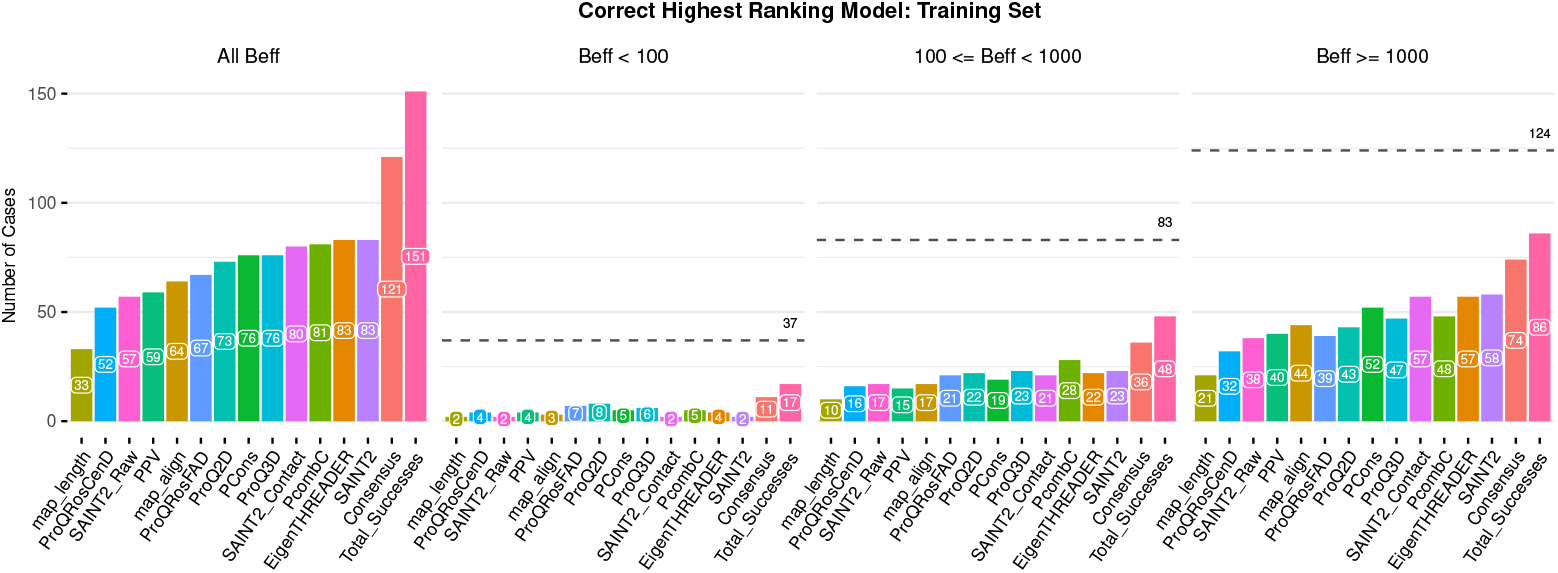
Number of targets out of the 244 targets in our Training set for which a correct model was produced and selected as the highest-ranked model according to 13 methods. Three SAINT2 scores (SAINT2, SAINT2_Contact and SAINT2_Raw), seven existing quality assessment scores (ProQ3D, ProQRosCenD, ProQRosFAD, Pcons, PcombC, ProQ2D and PPV), and three predicted contact map alignment scores (EigenTHREADER, Map align and map length) are shown, as well as all methods combined (“Consensus”) and the total number of targets with a correct model (“Total Successes”), for three *B*_eff_ bins and across all bins. The total number of targets in each *B*_eff_ bin is indicated with a dashed line.

We consider modelling to be a success if at least one correct model is produced for a target. For *B*_eff_ ≥ 1,000, SAINT2 produced correct models for 86 out of 124 targets (“Total Successes” in Fig 1). The two best methods for selecting correct models in this *B*_eff_ bin were the SAINT2 score and EigenTHREADER’s predicted contact map score; the highest-ranking models of these methods were correct (TM-score≥ 0.5) for 58 and 57 targets, respectively. The predicted contact potential of the SAINT2 score, SAINT2_Contact, also identified correct models for 57 targets, while only 38 were identified when this potential is excluded (SAINT2_Raw). Within this *B*_eff_ bin, the length of the map_align predicted contact map alignment selected correct models for the smallest number of targets, followed by PPV, the proportion of predicted contacts satisfied in the model. ProQRosCenD, a score based on the centroid knowledge-based energy potential Rosetta Centroid, also identified fewer correct models than the other scores, with a similar performance to SAINT2_Raw.

When considering 100 ≤ *B*_eff_ < 1,000, SAINT2 produced correct models for 48 out of 83 targets (“Total Successes” in Fig 1). PcombC performed the best at identifying correct models for this *B*_eff_ bin, with correct highest-ranking models for 28 targets, followed by the SAINT2 score and ProQ3D, each with 23 correct highest-ranking models. For *B*_eff_ < 100, SAINT2 produced correct models for 17 out of 37 targets. For these targets ProQ2D was the most successful, selecting a correct model for eight targets. Similar results were observed when considering the models output by SAINT2 for the Validation set (SI Fig 9).

As expected, these results demonstrate that methods using predicted contact information perform well on targets with more sequence data available, while knowledge-based scores are more informative for targets with less of this data. Overall, the SAINT2 score and EigenTHREADER identified correct highest-ranking models for 83 targets each, more targets than any other method (Fig 1). The best three methods, SAINT2, EigenTHREADER and PcombC identify correct models for different targets. Of the targets correctly identified by EigenTHREADER, 12 are not identified using SAINT2, and 17 are not identified by PcombC (SI Fig 10A). Incorporating EigenTHREADER scores when ranking the models produced by SAINT2 may therefore improve our ability to identify correct models. While these three methods are the major contributors (SI Fig 10B), we included all 12 methods into our random forest classifier as all methods had some predictive power.

### RFQAmodel: model quality assessment

Among the Validation set of 244 targets, 142 have a correct model within the 500 models produced by SAINT2. Selecting the highest-ranking model according to the SAINT2 score results in a correct model for 86 targets in this set. However, as the SAINT2 score cannot easily be compared between targets, it is difficult to infer for which targets the highest-ranked models are correct. We have trained a classifier, RFQAmodel, that assesses each model produced for a target and outputs a score, between 0 and 1, that the model has the correct fold.

We assessed the performance of RFQAmodel on our Validation set. Using a Receiver Operating Characteristic (ROC) curve, RFQAmodel achieved an area under the curve (AUC) of 0.95 for classifying all models for all targets as correct or incorrect, higher than all the individual component scores, including the best individual quality assessment score, Pcons (0.91), EigenTHREADER (0.84), and the SAINT2 score (0.77), as well as the other quality assessment scores ProQ2D (0.90), ProQ3D (0.89), ProQRosFAD (0.88), and PcombC (0.79) (SI Fig 11). In practice, we are interested in the classification of the highest-ranked model per target as correct or incorrect; for this task, RFQAmodel also outperforms the component methods (Fig 2 and SI Fig 11B).

**Fig 2.**
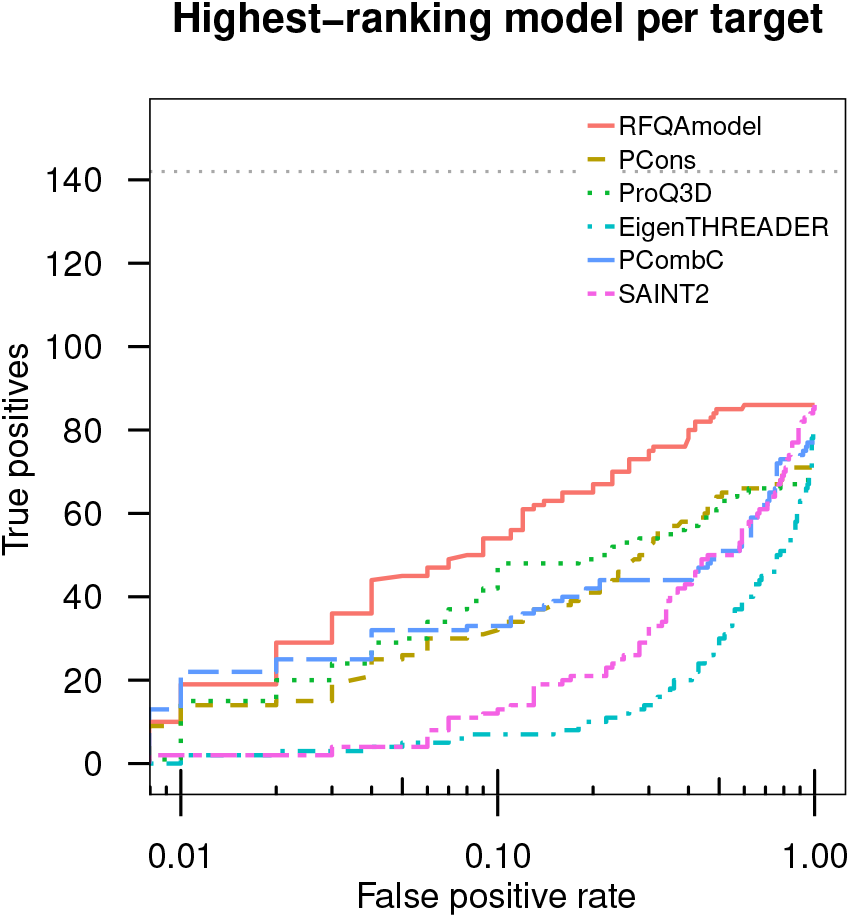
Classification of Validation Set targets. The number of targets with a correct highest-ranking model (true positives, TM-score ≥ 0.5) plotted against the false positive rate on a logarithmic scale, for the 244 targets in our Validation set. Curves are shown for the six highest-performing methods in Fig 1; curves for all component methods are shown in SI Fig 11. The grey dotted line indicates the total number of targets that had at least one correct model.

We divided the score output by RFQAmodel into four broad categories based on the Training set data: correct with high (>0.5), medium (between 0.3 and 0.5), or low (between 0.1 and 0.3) confidence, or predicted modelling failures (≤0.1) (SI Fig 12).

The models for a given target were ranked according to the RFQAmodel score, and targets were categorised based on the RFQAmodel score of the highest-ranking model. For each level of confidence, we assess whether the highest-ranking model (Top1) or the best of the top five highest-ranking models (Top5) is correct (Fig 3).

**Fig 3.**
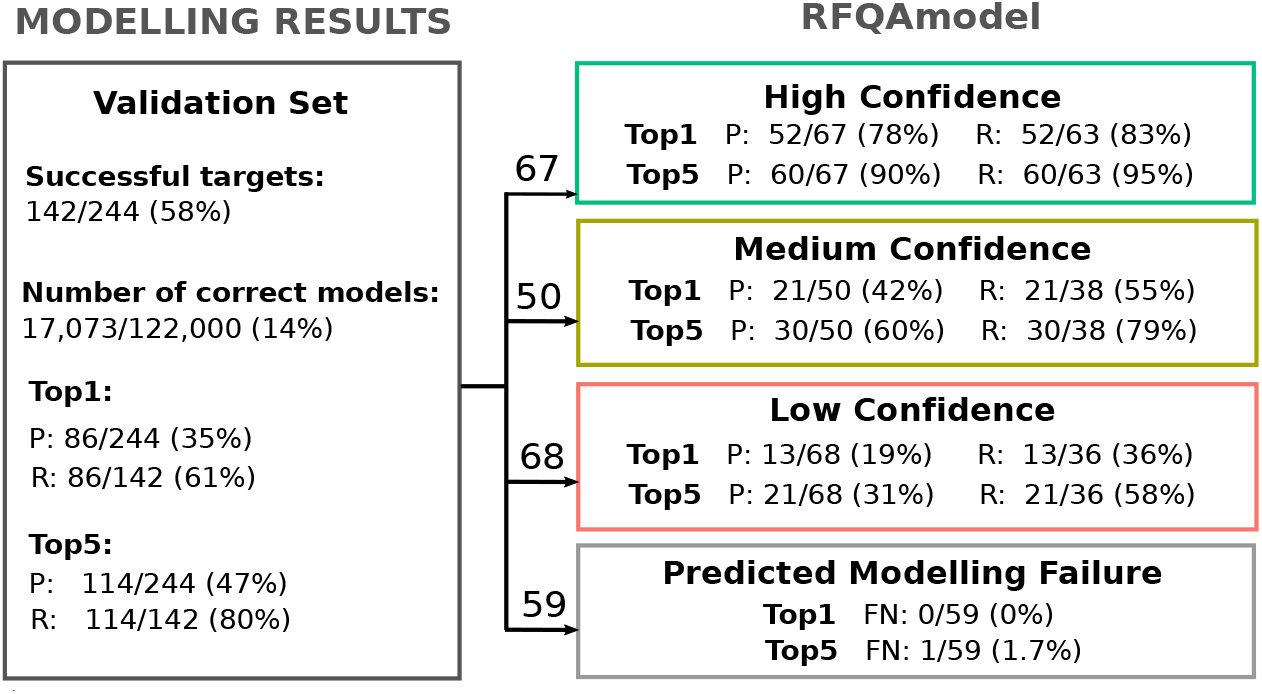
An overview of our classification protocol for the 244 modelling targets in our Validation set. The results of modelling (left, Section 3.2) and model quality assessment using RFQAmodel (right, Section 3.3) are shown. Modelling is considered successful for a given target if at least one model is correct (TM-score ≥ 0.5). For modelling results, models are ranked according to the SAINT2 score. For RFQAmodel results, models are ranked according to the RFQAmodel score. The precision (P) and recall (R) of the highest-ranked model (Top1) and the best of the top five highest-ranked models (Top5) are shown. For predicted modelling failures, the number of false negatives (FN) are shown.

When the models for each target in the Validation set are ranked according to the RFQAmodel score, the highest-ranking (Top1) model is correct for 86 of 244 targets. This is exactly the same as the number of correct highest-ranking models when ranked according to SAINT2; the difference is that RFQAmodel assigns a likelihood that each model is correct. RFQAmodel predicts that modelling has failed (≤0.1) for all models for 59 targets. For 5 of these targets there was at least one correct model in the 500, but the highest-ranked model was not correct for any. Excluding these 59 targets reduces our Validation set from 244 to 185 targets, of which 137 have a correct model.

The highest-ranking (Top1) model was predicted to be correct with low confidence for 68 targets. This model was correct for 13 of these targets (19% precision), and 21 targets had a correct model in the top five (Top5) highest-ranking models (31% precision).

The highest-ranking model was predicted to be correct with medium confidence for 50 targets. The highest-ranked model was correct for 21 of these targets (42% precision), and the best out of the top five highest-ranking models was correct for 30 targets (60% precision).

The highest-ranking model was predicted to be correct with high confidence for 67 targets. This model was correct for 52 out of these 67 high-confidence targets (78% precision), and the best out of the top five highest-ranking models was correct for 60 of these targets (90% precision).

When considering the combined results for the 117 targets with highest-ranking models predicted to be correct with high or medium confidence, this model was correct for 73 targets (62% precision), and the best out of the top five highest-ranking models was correct for 90 of these targets (77% precision).

### Comparison to methods used in large-scale studies

We compared RFQAmodel to two methods that have been used to evaluate the success of large-scale predictions of unknown protein structures by Michel et al. [12] and Ovchinnikov et al. [10]. In the study by Michel et al., the authors used the PcombC score cut-off that achieved a false positive rate (FPR) of 0.01 and 0.1 on the benchmarking set to predict whether models were correct (TM-score ≥ 0.5) [12]. PcombC is one of the scores used in RFQAmodel, so it is unsurprising that RFQAmodel is able to achieve better performance (Fig 2). Compared to PcombC, RFQAmodel performs similarly at an FPR of 0.01, but identifies a correct model for more targets at 0.1 (see Fig 2).

To compare RFQAmodel with the method used in Ovchinnikov et al., we calculated the mean pairwise TM-score of the 10 models with the highest ProQ3RosCenD score out of the 500 models generated for each target, and classified targets above 0.65 as correct [10]. This method classified 21 targets as correct, of which 19 had a correct highest-ranking model. A similarly high precision was achieved using ProQ3RosFAD instead of ProQ3RosCenD (19 out of 22). Using RFQAmodel, a similar precision with higher recall can be achieved with a cut-off of 0.7, with 26 of 29 targets having a correct highest-ranking model (Fig 4, solid lines). Using the high confidence cut-off for RFQAmodel we achieve 78% precision and 37% recall. At this level of recall, the ProQRosCenD method achieves a precision of 36% (Fig 4, dashed lines). The difference between the methods appears to be the ability of RFQAmodel to identify correctly modelled targets with fewer correct models (Fig 4).

**Fig 4.**
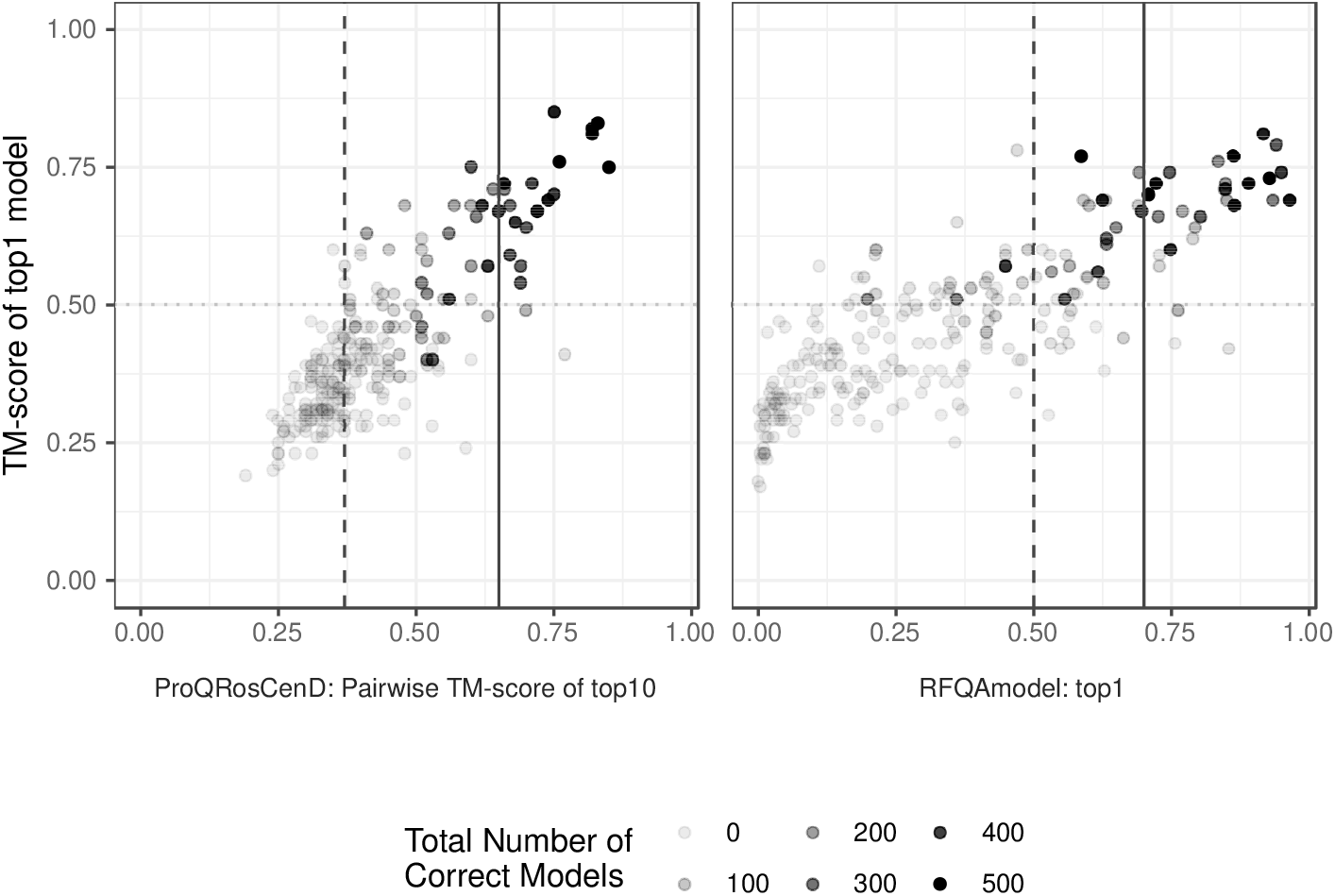
Using convergence or RFQAmodel to identify correct models. The TM-score of the highest-ranking model for each of the 244 targets in the Validation set according to ProQRosCenD and RFQAmodel, against the mean pairwise TM-score of the 10 highest-ranking models (ProQRosCenD, left) or the score of the highest-ranking model (RFQAmodel, right). Targets with a mean pairwise TM-score greater than 0.65 are predicted to be correct (solid line, left); a similar precision is achieved with an RFQAmodel cut-off of 0.7 (solid line, right). A pairwise TM-score cut-off of 0.37 (dashed line, left) achieves a similar recall to the high confidence cut-off of RFQAmodel (dashed line, right). Targets for which fewer correct models were generated among the 500 models are shown with lighter circles.

### CASP12 and CASP13 Quality Assessment

RFQAmodel was trained and validated on models generated using SAINT2. In order to test its performance on models generated by other methods, we used RFQAmodel to classify models for the 57 CASP12 and 72 CASP13 Quality Assessment targets (see Methods). We used the stage2 set: the 150 highest-ranking models per target selected from the server predictions, with up to five models contributed by 93 different methods. The targets are not divided into constituent domains for the evaluation of quality assessment methods in CASP. As RFQAmodel is designed to assess the output of template-free protein structure prediction protocols as correct or incorrect, here, we only evaluate its performance on the 33 CASP12 and 34 CASP13 targets containing domains classified as free-modelling targets. RFQAmodel performs well on models of the easier template-based modelling targets, which tend to be globally more accurate (SI Table 4).

We used RFQAmodel, trained on the SAINT2 Training set, to classify models in the CASP12 and CASP13 sets as either correct or incorrect. Of the 67 free-modelling targets, 47 targets had at least one correct model. When classified using RFQAmodel, 31 targets had a high confidence highest-ranking model, of which 21 were correct (68% precision, 31% recall).

To assess the performance against other quality assessment techniques, we compared RFQAmodel to the predictions submitted to CASP13 for free-modelling targets. These blind predictions were submitted between May and July 2018, and made publically available in December 2018. We find that RFQAmodel performs similarly to the top performing methods at classifying individual models and the highest-ranking model as correct or incorrect (Fig 5).

**Fig 5.**
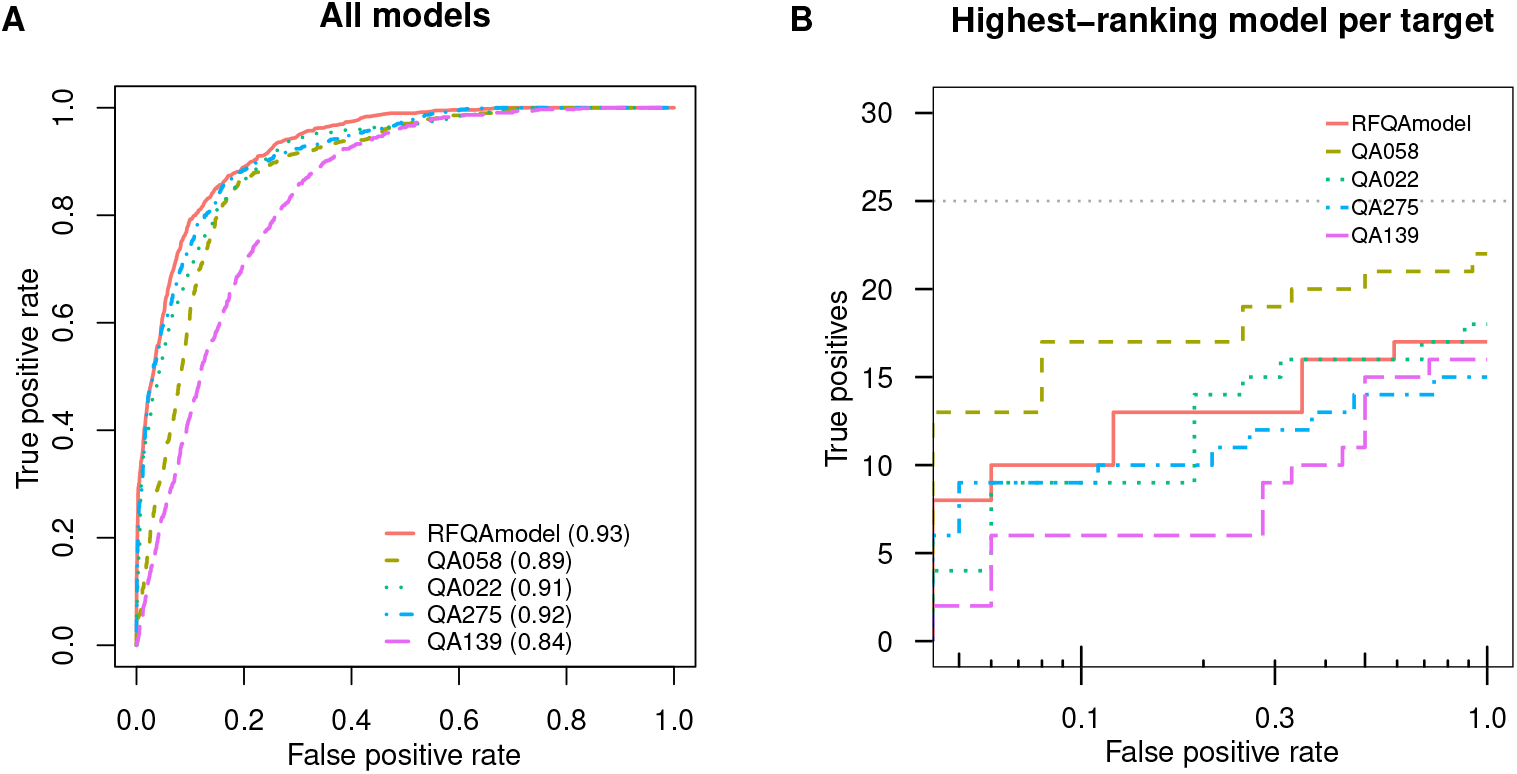
Classification of CASP13 free-modelling targets. Receiver Operating Characteristic (ROC) Curves for the classification of all models into whether they were correct (TM-score ≥ 0.5) or incorrect according to RFQAmodel and four quality assessment scores submitted for the 34 free-modelling targets in the CASP13 set. The area under the ROC curve (AUC) for each method is shown in brackets. B) The number of targets with a correct highest-ranking model (true positives) plotted against the false positive rate on a logarithmic scale. The grey dotted line indicates the total number of targets that had at least one correct model.

### Iterative model generation and quality assessment

The optimal number of models to generate using SAINT2 is 10,000, but RFQAmodel may enable us to focus our computational efforts more efficiently by identifying the targets for which fewer models are sufficient to generate good models. It may be possible to improve modelling results by iteratively generating more models for the predicted modelling failures and applying RFQAmodel until modelling it predicted to have succeeded with the required confidence.

In order to assess this application, we chose five targets for which RFQAmodel predicted the highest-ranking model to be correct with low confidence or modelling failures based on the initial 500 models. We then iteratively generated 10,000 models in intervals of 500 models; at each interval we reassessed the model ensemble and compared the TM-score of the best of the top5 highest-ranking models (Fig 6). As generating and assessing 10,000 models is computationally expensive, carrying out this analysis on all 244 targets in the Validation set is infeasible.

**Fig 6.**
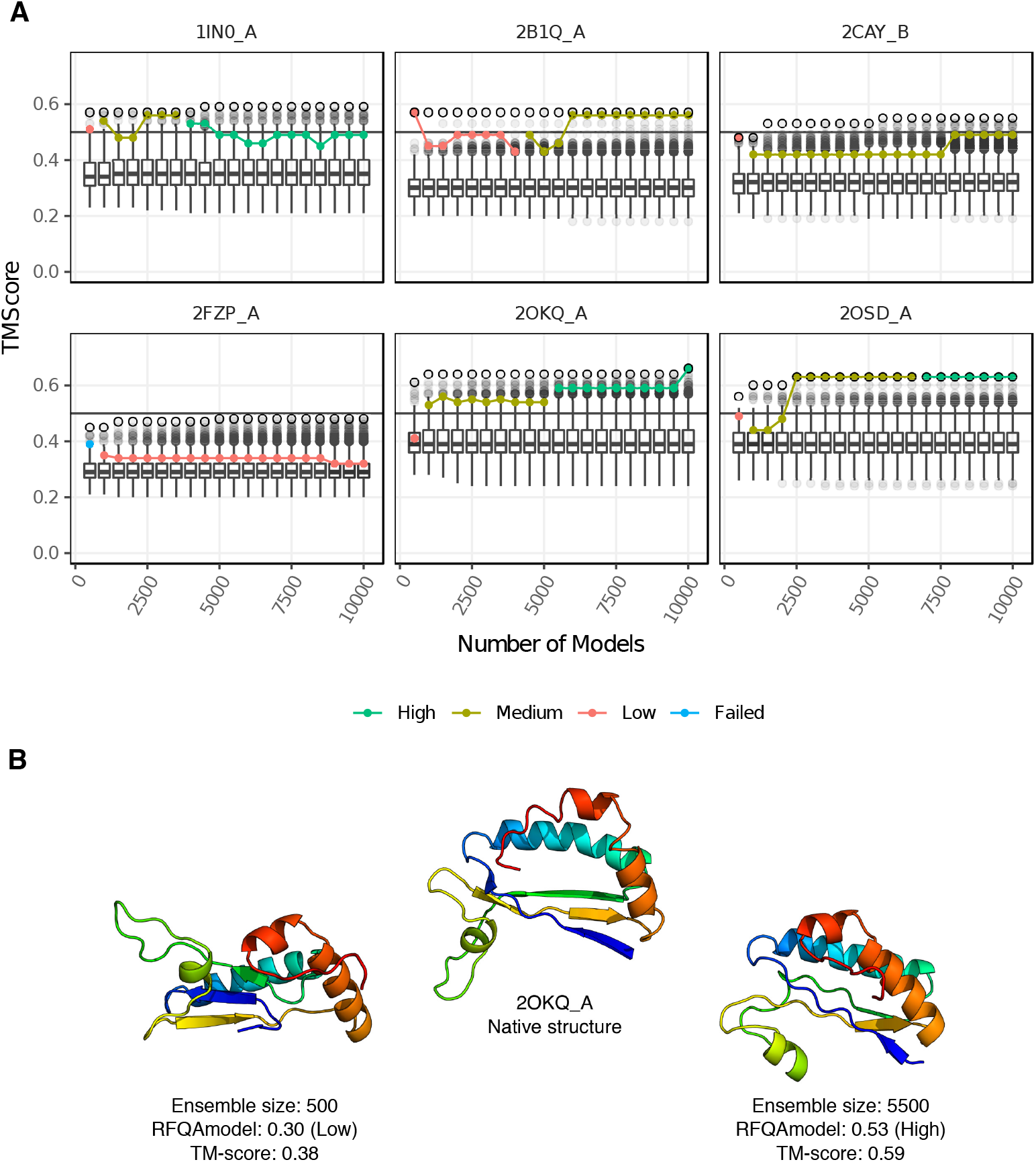
RFQAmodel classification improves with ensemble size. A) Six targets that were initially classified by RFQAmodel as low confidence or failed were chosen. The TM-scores of the models are shown in boxplots, as the number of models generated for each target is increased in increments of 500 from 500 to 10,000. The best model (highest TM-score) is highlighted with a black circle. The TM-score of the best of the top5 highest-ranking model according to RFQAmodel for each ensemble size is indicated with a filled circle, coloured according to the Confidence. B) The native structure of 2OKQA (centre) compared to the highest-ranked model according to RFQAmodel after 500 models were generated (left) and after 5,500 models were generated (right), at which point a high confidence RFQAmodel score was achieved.

For one target, 2FZPA, no correct models were generated, and RFQAmodel classified the highest-ranking model as failed or low confidence for all ensemble sizes. For another target, 2CAYB, the confidence increased from low to medium confidence, but a correct model was never identified. For 1IN0A, a high-confidence model was identified once the ensemble size reached 4,000, and this model was correct. Interestingly, if model generation continues, the quality of the highest-ranking models decreases after 6,000 models. For the remaining three targets, RFQAmodel selected better models with higher confidence as the ensemble size increased (2B1QA, 2OKQA and 2OSDA). For example, for 2OKQA the highest-ranked model of the initial 500 models had a low-confidence RFQAmodel score of 0.3 (TM-score 0.38). After 1,000 models were generated, the highest-ranked model had a medium-confidence score of 0.44 (TM-score 0.53). Once the ensemble size reaches 5,500 the highest-ranked model had a high-confidence RFQAmodel score of 0.53, and a TM-score of 0.59 (Fig 6B). These results demonstrate how RFQAmodel could be used to guide computational efforts and thus and increase the number of targets for which we have a good predicted structure.

## Discussion

We show, as have others, that both modelling and quality assessment are more likely to succeed for targets that are shorter, mostly alpha-helical, or have higher *B*_eff_ values (e.g. SI Fig 7) [1, 8, 20]. Previous attempts at estimating quality assessment success have used training and test sets that were not balanced in length and number of effective sequences (e.g. [12]), which may result in inconsistent performance when applied to other sets. In order to ensure as accurate an estimate of performance as possible, we designed our Training and Validation sets to be well-balanced in terms of these features.

Using our Training set we built RFQAmodel, which uses the contact map alignment scores EigenTHREADER and map align in addition to existing quality assessment scores to estimate model quality. For targets with sufficient sequence information, we found that EigenTHREADER identifies correct models for more targets than a number of existing single-model, consensus, and hybrid model quality scores (Fig 1). Eight of these targets were not captured by the two other top performing methods, SAINT2 and PcombC. This indicates that predicted contact map alignment scores are, at least to some extent, orthogonal to existing model quality assessment scores.

Unlike many existing quality assessment scores, RFQAmodel was designed to output a score that indicates the likelihood that a model is correct. On our Validation set it identifies, with high confidence, a single correct model for 67 of 244 targets with 78% precision. RFQAmodel outperformed the component quality assessment methods, in agreement with previous studies where combining methods improves performance [4, 12, 21]. When compared to methods used to identify successfully modelled targets in large-scale protein structure prediction studies [10, 12], RFQAmodel achieved a higher recall and was able to identify successfully modelled targets with fewer correct models in their ensemble. This suggests that by using RFQAmodel it may be possible to identify more modelling successes in large-scale studies.

While RFQAmodel was developed and trained using our template-free protein structure prediction protocol, SAINT2, we assessed its suitability for use with other protocols. We tested RFQAmodel on ensembles of models from a large number of different protocols for 56 CASP12 and CASP13 free-modelling targets. RFQAmodel classified the highest-ranking model as correct with high confidence for 38% of targets with 81% precision and 85% recall. While this demonstrates that RFQAmodel can be used to classify models generated by methods other than SAINT2, the performance of RFQAmodel may be improved by training on models from a variety of other protocols.

RFQAmodel was not trained for other quality assessment tasks, such as predicting the absolute quality of models. Furthermore, unlike some methods (including ProQ3D and PCons), RFQAmodel does not estimate the local (per-residue) quality of models. However, we found that it performed comparably to the top-performing methods in CASP13 at selecting a correct model for each target.

Finally, our protocol is able to reduce the computational cost of protein structure prediction, which is a common limitation for large-scale studies. The assignment of confidence enables us to identify the targets for which 500 models are sufficient to generate good models with high confidence. We can then iteratively generate more models for the medium, low confidence, or failed targets and apply RFQAmodel until modelling is predicted to have succeeded with high confidence, focussing computational efforts more efficiently.

## Supporting information

**SI Table 1 Properties of the 8,005 protein chains representing each of the Pfam domains mapped to PDB structures**.

**SI Table 2 Properties of the 4,728 protein chains with SCOPe annotations chosen to represent unique Pfam families mapped to PDB structures**.

**SI Table 3 Properties of the 488 protein domains chosen to comprise our Training and Validation data sets**.

**SI Table 4 RFQAmodel performance for all CASP12 and CASP13 free-modelling and template-based modelling targets**.

**SI Section 1 RFQAmodel Random Seed**.

**SI Section 2 Data sets & culling process**.

**SI Section 3 Number of effective sequences definition**.

**SI Fig 1 SCOP classes of representative chains**.

**SI Fig 2 Domain lengths and resolutions of Training and Validation sets**.

**SI Section 4 Prediction of sequence-based descriptors**.

**SI Section 5 Estimating the number of models required**.

**SI Section 6 Modelling results**.

**SI Fig 7 Modelling success rate by *B*_eff_, SCOP class, and domain length**.

**SI Fig 8 Modelling success rate by both *B*_eff_ and domain length**.

**SI Fig 9 Ranking of models for Validation set targets**.

**SI Fig 10 The number of targets in our Training set for which the highest-ranking models are correct according to the three overall best methods and all methods combined**.

**SI Fig 11 Classification of Validation set targets**.

**SI Fig 12 Confidence categorisation of RFQAmodel scores**.

## Acknowledgments

The authors would like to acknowledge the Oxford Protein Informatics Group and Dr Sebastian Kelm for their intellectual input and comments on the draft.

